# Grass Carp Reovirus (GCRV) Giving Its All to Suppress IFN Production by Countering MAVS-TBK1 Activation

**DOI:** 10.1101/792408

**Authors:** Long-Feng Lu, Zhuo-Cong Li, Can Zhang, Xiao-Yu Zhou, Yu Zhou, Jing-Yu Jiang, Dan-Dan Chen, Shun Li, Yong-An Zhang

## Abstract

As a crucial signaling pathway for interferon (IFN) production, the RIG-I-like receptor (RLR) axis is usually the host target of viruses to enhance viral infection. To date, though immune evasion methods to contrapose IFN production have been characterized for a series of terrestrial viruses, the strategies employed by fish viruses remain unclear. Here, we report that all grass carp reovirus (GCRV) proteins encoded by segments S1 to S11 interact with fish RLR factors, specifically for mitochondrial antiviral signaling protein-TANK-binding kinase 1 (MAVS-TBK1) signaling transduction, leading to decreased IFN expression. First, the GCRV viral proteins blunted the MAVS-induced expression of IFN but had little effect on TBK1-induced IFN expression. Subsequently, interestingly, co-immunoprecipitation experiments demonstrated that all GCRV viral proteins interacted with several RLR cascades, especially with TBK1. To further illustrate the mechanisms of these interactions between GCRV viral proteins and host RLRs, two of the viral proteins, NS79 (S4) and VP3 (S3), were selected as representative proteins for the study. The obtained data demonstrated that NS79 did not affect the stability of the host RLR protein, but was phosphorylated by gcTBK1, leading to the reduction of host substrate gcIRF3/7 phosphorylation. On the other hand, VP3 degraded gcMAVS and the degradation was significantly reversed by 3-MA. The biological effects of both NS79 and VP3 were consistently found to be related to the suppression of IFN expression and the promotion of viral evasion. Our findings shed light on the special evasion mechanism utilized by fish virus through IFN regulation, which might differ between fish and mammals.

**Author summary:** The RLR signaling pathway is crucial for IFN induction when host cells are infected with virus and RLR factors are targeted by virus. To date, the evasion mechanisms of fish viruses remain mysterious. In this study, we reveal that all 11 GCRV proteins interact with fish RLR factors and suppress the activation of MAVS-TBK1 signaling transduction, leading to the reduction of IFN expression. Two viral proteins were employed as examples to investigate the different evasion mechanisms of GCRV. These findings reveal the novel countermeasures used by fish virus to avoid the host IFN response.

## Introduction

Interferons (IFNs) are considered the first and fundamental line of defense against viral invasion in both mammals and fish [1, 2]. The production of IFNs is triggered by signal transduction once the host cell senses viral components [3]. For most viruses, the virion is composed of viral nucleic acids, such as DNA, double stranded RNA (dsRNA), single stranded RNA (ssRNA), and surface glycoproteins [4]. Pathogen-associated molecular patterns (PAMPs) (including viral nucleic acids and proteins) are usually recognized by pattern recognition receptors (PRRs), which are expressed on the surface and cytoplasm of host cells [4]. Among the PRR members, the retinoic acid inducible gene-I (RIG-I) mediates a pivotal signaling pathway termed the RIG-I-like receptor (RLR) pathway, which significantly activates IFN transcription [5]. Upon binding with the viral RNA, RIG-I or melanoma differentiation-associated gene 5 (MDA5) recruits the downstream adaptor mitochondrial antiviral signaling protein (MAVS, also called VISA, IPS-1, or Cardif) [6-9] and the mediator of IFN regulatory factor 3 (IRF3) activation (MITA, also termed STING, MPYS, or ERIS) [10-13], then activates TANK-binding kinase 1 (TBK1). Activated TBK1 further phosphorylates IFN regulatory factor 3/7 (IRF3/7), triggering their dimerization and nuclear translocation to bind to IFN stimulation response elements (ISREs) and initiate the transcription of IFN [14-16].

MAVS is essential for host innate immune responses against viral infection [8]. It contains an N-terminal caspase recruitment domain (CARD), a middle proline-rich domain, and a C-terminal transmembrane (TM) domain [17, 18]. MAVS is an adaptor protein involved in virus-triggered IFN signaling and regulates virus-induced apoptosis to limit viral replication [19]. In fish, multiple-sequence alignments and phylogenetic analysis have demonstrated that teleost fish possess a *mavs* gene that is involved in the regulation of IFN production [20]. For instance, in zebrafish (*Danio rerio*), MAVS overexpression results in a robust activation and upregulation of IFN and IFN-stimulated genes (ISGs) in response to RNA and DNA virus infection [21].

TBK1 is a non-canonical IκB kinase (IKK) that plays critical roles in IFN induction and innate antiviral immunity [22]. It consists of three domains: an N-terminal serine/threonine kinase domain (KD), a ubiquitin-like domain (ULD), and a C-terminal domain (CTD) (also known as two C-terminal coiled coil domains) [23]. Actually, as a ubiquitously expressed kinase, besides the TBK1-IRF3/7 pathway, TBK1 participates in several other signaling pathways such as autophagy and cell cycle control [24, 25]. TBK1 has been characterized in many fish species, including zebrafish, crucian carp (*Carassius aumtus*), and grass carp (*Ctenopharyngodon idella*) [26-28]. Overexpression of grass carp TBK1 induces the upregulation of IFN1 upon grass carp reovirus (GCRV) infection [26].

Viruses have evolved elaborate strategies to evade or abrogate the host IFN signaling pathway for their replication. As MAVS and TBK1 are key molecules in the RLR pathway for the activation of IFN production, they are popular targets of viral antagonists. For example, the Newcastle disease virus (NDV) V protein inhibits IFN production through targeting MAVS for ubiquitin-mediated degradation via the E3 ubiquitin ligase RING-finger protein 5 (RNF5) [29]. Similarly, the 3C protein of coxsackievirus B3 (CVB3) and porcine reproductive and respiratory syndrome virus (PRRSV) suppresses IFN activation by cleaving MAVS [30, 31]. The NS3 protein of the hepatitis C virus (HCV) blocks IFN signaling by binding to TBK1 and disrupts the interaction between TBK1 and IRF3 [32]. Finally, the leader proteinase (Lb^pro^) of foot-and-mouth disease virus (FMDV) counteracts host antiviral responses via mediating TBK1 deubiquitination [33].

GCRV, a highly virulent pathogenic agent of fish, has caused severe epidemic outbreaks of hemorrhagic disease and resulted in tremendous mortality among grass carp [34]. It is a dsRNA virus that belongs to the genus *Aquareovirus* in the family *Reoviridae* [35]. Based on genomic and biological characteristics, known GCRV strains can be clustered into three groups (groups I-III), and studies have demonstrated that the highest mortality for grass carp is usually caused by group II GCRV [35]. The GCRV genome consists of 11 segments (termed S1–S11) and is encased in a multilayered icosahedral capsid shell; until now, the proteins encoded by these segments were unclear [36, 37]. In previous studies, fish IFNs and ISGs exhibited a powerful capacity to defend against infection by GCRV [26, 38, 39]. However, GCRV leads to outbreaks of fish hemorrhagic disease [34], indicating that the virus employs a strategy to evade the host IFN response for successful infection. In our previous study, the GCRV VP41 inhibited MITA phosphorylation by acting as a decoy substrate of TBK1, thus reducing IFN production and facilitating viral replication. However, GCRV could possess a number of different strategies to elude host defense mechanisms. Therefore, uncovering the other mechanisms used by GCRV to inhibit the activation of IFN signaling is warranted.

In this study, we show that all GCRV viral proteins encoded by S1 to S11 associate with fish RLR factors, specifically blocking the MAVS-induced IFN expression. Using NS79 and VP3 as representative proteins, we found that NS79 reduces gcMITA phosphorylation by acting as a decoy substrate of gcTBK1 while VP3 degrades gcMAVS in an autophagosome-dependent manner, ultimately blocking IFN production and facilitating virus replication. These results uncovered two distinct evasion strategies used by GCRV to escape the host IFN system by targeting gcRLR factors. Our findings will lay the foundation for further study of the crosstalk between the host IFN response and viral infection in fish species.

## Results

### The MAVS-TBK1 pathway for IFN activation is blocked by all viral proteins of GCRV

Unlike in mammals, there are multiple types of IFNs in fish. The current study focuses on the grass carp. There are four homologs of type I IFN in grass carp, termed gcIFN1-gcIFN4. To date, the role of these members under influence remain unclear; therefore, the characterization of these IFNs was first performed. Treatment with poly I:C, a mimic of viral RNAs, resulted in a significant increase in gcIFN1 promoter (gcIFN1pro) activity compared with other IFNs (Fig 1A). As fish RLR cascades are pivotal IFN activators, the upstream factors gcRIG-I, gcMAVS, and gcTBK1 were employed for IFN identification. Consistent with the above result, gcIFN1 displayed remarkable activation under gcRLR stimulation (Fig 1B-1D). Therefore, gcIFN1 was selected as the reporter gene for subsequent assays.

**Fig 1.**
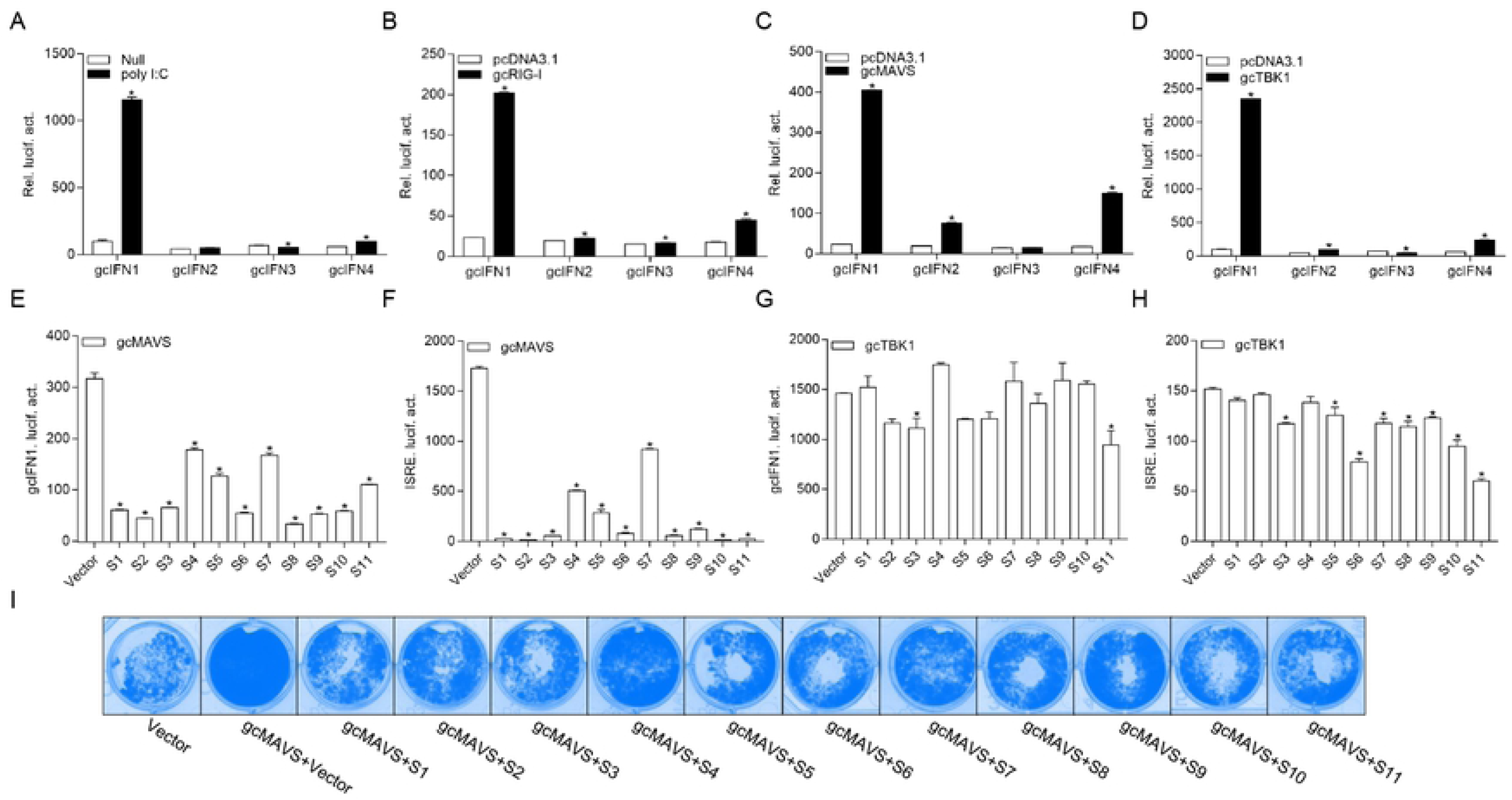
GCRV viral proteins encoded by S1-S11 suppress the MAVS-mediated activation of IFN1 and its antiviral effect. (A) Induction of gcIFN1, gcIFN2, gcIFN3, gcIFN4 promoters by poly (I:C). GCO cells were seeded in 24-well plates overnight were transfected with 0.25 μg gcIFN1pro-Luc, gcIFN2pro-Luc, gcIFN3pro-Luc, or gcIFN4pro-Luc, 50 ng pRL-TK was used as an internal control. At 24 h post-transfection, cells were transfected with poly (I:C) (1 μg/ml) or left untreated (null). The luciferase assay was performed 24 h after stimulation. (B-D) Activation of gcIFN1, gcIFN2, gcIFN3, gcIFN4 by overexpression of gcRIG-I or gcMAVS or gcTBK1. GCO cells seeded in 24-well plates overnight were co-transfected with pcDNA3.1-gcRIG-I (B) or pcDNA3.1-gcMAVS (C) or pcDNA3.1-gcTBK1 (D) or empty vector and gcIFN1pro-Luc, gcIFN2pro-Luc, gcIFN3pro-Luc or gcIFN4pro-Luc at the ratio of 1:1. pRL-TK was used as a control. Luciferase activities were analyzed at 24 h post-transfection. (E and F) GCRV viral proteins inhibit gcIFN1 and ISRE activation mediated by gcMAVS. GCO cells were seeded in 24-well plates and co-transfected with pcDNA3.1-gcMAVS and empty vector or pcDNA3.1-S1, or pcDNA3.1-S2, or pcDNA3.1-S3, or pcDNA3.1-S4, or pcDNA3.1-S5, or pcDNA3.1-S6, or pcDNA3.1-S7, or pcDNA3.1-S8, or pcDNA3.1-S9, or pcDNA3.1-S10, or pcDNA3.1-S11, plus gcIFN1pro-Luc (E) or ISRE-Luc (F) at the ratio of 1:1:1. pRL-TK was used as a control. At 24 h post-transfection, cells were lysed for luciferase activity detection. (G and H) GCRV viral proteins have no apparent effects on gcTBK1-mediated activation of gcIFN1 (G) and ISRE reporters (H). GCO cells were seeded in 24-well plates and co-transfected with pcDNA3.1-gcTBK1 and empty vector or pcDNA3.1-S1, or pcDNA3.1-S2, or pcDNA3.1-S3, or pcDNA3.1-S4, or pcDNA3.1-S5, or pcDNA3.1-S6, or pcDNA3.1-S7, or pcDNA3.1-S8, or pcDNA3.1-S9, or pcDNA3.1-S10, or pcDNA3.1-S11, plus gcIFN1pro-Luc (G) or ISRE-Luc (H) at the ratio of 1:1:1. pRL-TK was used as a control. At 24 h post-transfection, cells were lysed for luciferase activity detection. The promoter activity is presented as relative light units normalized to Renilla luciferase activity. Data were expressed as mean ± SEM, *n* = 3. Asterisks indicate significant differences from control (*, *p* < 0.05). (I) The S1-S11 segments inhibit the antiviral effect of gcMAVS. EPC cells were seeded in 24-well plates overnight and transfected with indicated plasmids for 24 h, then the cells were infected with SVCV (MOI = 0.001), at 48 h post infection, the cells were fixed with 4% PFA and stained with 1% crystal violet. All experiments were repeated for at least three times with similar result.

The 11 segments of GCRV were subcloned into eukaryotic expression vectors to investigate the roles of GCRV viral proteins in IFN regulation. The GCRV S1-S11 and gcRLR factors were co-expressed and the activation of gcIFN1pro was monitored. ISRE is considered a transcription factor-binding motif in the promoter regions of IFNs and ISGs, facilitating gene transcription. Interestingly, as gcMAVS is the pivotal upstream factor of gcTBK1, all 11 viral proteins suppressed its effects on gcIFN1 and ISRE activity induction (Fig 1E and 1F). Compared with the impressive antagonism against gcMAVS activation, the regulation of gcTBK1 by the GCRV viral proteins was remarkably weaker (Fig 1G and 1H). The *Danio rerio* (Dr) MAVS, TBK1, and IFNφ1pro were also employed to assess the conservation of MAVS regulation by the GCRV viral proteins. Similar to the results in grass carp, the activation of DrIFNφ1pro and ISRE induced by DrMAVS was reduced by S1-S11 (S1A and S1B Fig), while DrTBK1-induced DrIFNφ1pro and ISRE activities were unaffected by GCRV viral proteins (S1C and S1D Fig). Next, we examined whether GCRV viral proteins had an effect on the gcMAVS-mediated antiviral response. We transfected EPC cells with gcMAVS, together with a control plasmid or S1-S11 expression plasmids. The transfected cells were infected with SVCV. As shown in Fig 1I, stronger CPEs were observed in the GCRV viral proteins groups at 2 d post-infection. These results demonstrate that the GCRV viral proteins impaired MAVS-induced activation of IFN and inhibited the MAVS-mediated antiviral response, but had little effect on TBK1, indicating that they blocked the signaling transduction from MAVS to TBK1.

### TBK1 is the common target of GCRV viral proteins

Combined with the observation that MAVS recruits TBK1 in the signaling transduction process, the above results suggest that the GCRV viral proteins might target the host TBK1. Co-IP experiments were performed to characterize the relationship between the GCRV viral proteins and gcTBK1. When Myc-tagged GCRV S1 to S11 and Flag-tagged gcTBK1 were overexpressed, the anti-Myc Ab-immunoprecipitated S1 to S11 protein complexes were recognized by the anti-Flag Ab (Fig 2A), and vice versa (Fig 2B-2L), demonstrating that all GCRV viral proteins were associated with grass carp TBK1. We confirmed the ubiquitous association of all GCRV viral proteins with TBK1 in different fish species by also assaying the corresponding viral proteins with zebrafish RLR factors. The results showed that all GCRV viral proteins interacted with the RLR molecules, particularly with DrTBK1 (S2 Fig), indicating that the interaction of viral proteins with fish TBK1 is conserved. To our knowledge, these data manifest a novel mechanism that all GCRV viral proteins may associate with TBK1, which is the key factor in IFN activation.

**Fig 2.**
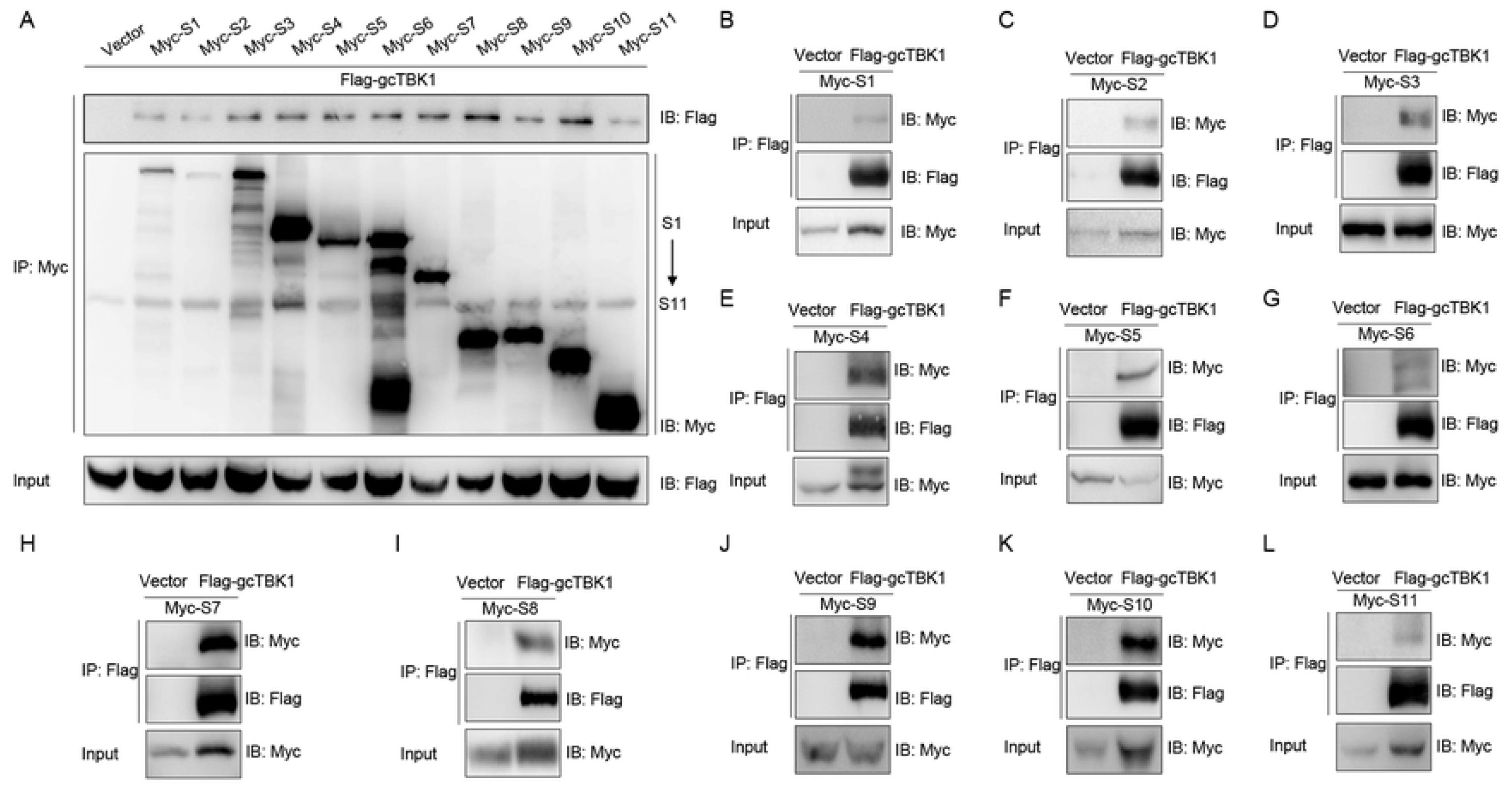
The interaction between GCRV viral proteins and gcTBK1. (A-L) HEK 293T cells seeded in 10-cm^2^ dishes were transfected with the indicated plasmids (5 μg each). After 24 h, cell lysates were immunoprecipitated (IP) with an anti-Myc/Flag affinity gel. Then the immunoprecipitates and cell lysates were analyzed by immunoblotting (IB) with the anti-Flag and anti-Myc Abs, respectively. All experiments were repeated at least three times, with similar results.

### GCRV NS79 or VP3 suppresses IFN induction and colocalizes with gcMAVS and gcTBK1

The above observations revealed that all GCRV viral proteins blunt MAVS-induced IFN activation. To further investigate the specific mechanisms underlying how GCRV proteins evade IFN responses, we chose S3 and S4-encoded proteins for the subsequent assays. The S3 and S4 segments of group II GCRV were predicted to encode the respective inner core protein VP3 and the nonstructural protein NS79, which are involved in viral inclusion body formation [40]. As shown in Fig 3A, poly I:C stimulation induced the activation of gcIFN1pro; however, the induction was significantly blocked by the overexpression of NS79 or VP3. NS79 or VP3 also suppressed ISRE activity upon transfection with poly I:C (Fig 3B). Fish RLR factors are crucial for triggering IFN expression. Consequently, ORFs of the genes involved in the *grass carp* RLR signaling pathway, including gcMAVS, gcTBK1, gcMITA, gcIRF3, and gcIRF7, were cloned according to the sequences in a public database. As shown in Fig 3C and 3D, overexpression of the gcRLR factors led to a significant induction of gcIFN1pro activity, and the activation of gcIFN1 induced by gcMAVS was inhibited by co-transfection with NS79 or VP3. However, ectopic expression of NS79 or VP3 did not affect the gcTBK1-stimulated gcIFN1pro activity. Similarly, the ISRE activity upregulated by gcMAVS but not gcTBK1 also decreased with the overexpression of NS79 or VP3 (Fig 3E and 3F). Given that gcTBK1 is downstream of gcMAVS, these results indicate that NS79 or VP3 likely blocked IFN production via the negative regulation of gcMAVS or gcTBK1.

**Fig 3.**
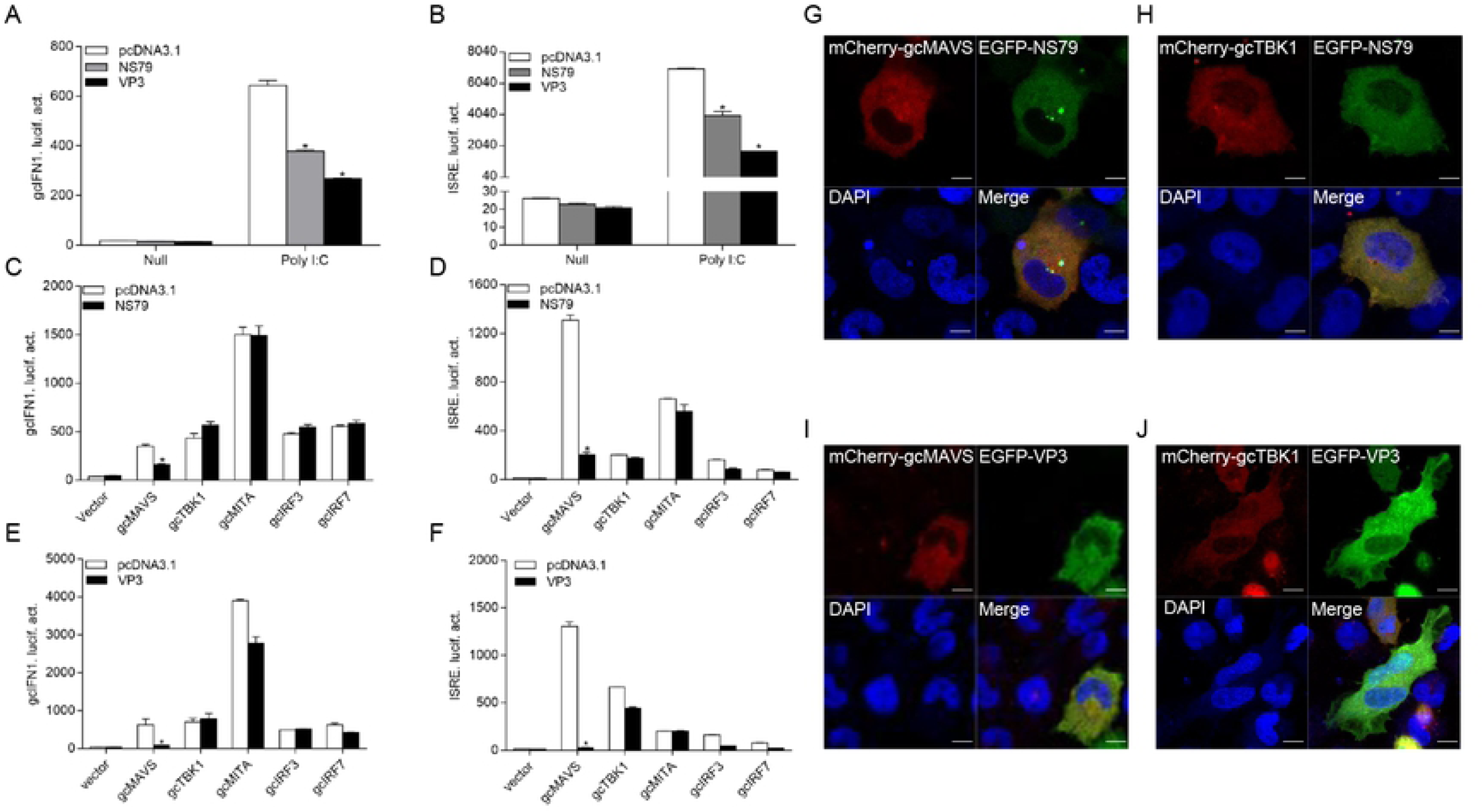
GCRV NS79 or VP3 inhibits gcIFN1 and ISRE activation and colocalizes with gcMAVS and gcTBK1. (A and B) Overexpression of NS79 or VP3 inhibits poly (I:C)-induced gcIFN1pro/ISRE activation. EPC cells were seeded in 24-well plates overnight and co-transfected with 0.25 μg gcIFN1pro-Luc (A) or ISRE-Luc (B), and 50 ng pRL-TK, plus 0.25 μg empty vector or pcDNA3.1-NS79, or pcDNA3.1-VP3. At 24 h post-transfection, cells were transfected with poly (I:C) (1 μg/ml) or left untreated (null). The luciferase assay was performed 24 h after stimulation. (C-F) NS79 or VP3 suppresses gcIFN1pro/ISRE activation mediated by gcRLRs. EPC cells were seeded in 24-well plates and co-transfected with gcRLRs-expressing plasmids and empty vector or pcDNA3.1-NS79, or pcDNA3.1-VP3, plus gcIFN1pro-Luc or ISRE-Luc at the ratio of 1:1:1. pRL-TK was used as a control. At 24 h post-transfection, cells were lysed for luciferase activity detection. The promoter activity is presented as relative light units normalized to Renilla luciferase activity. Data were expressed as mean ± SEM, *n* = 3. Asterisks indicate significant differences from control (*, *p* < 0.05). (G-J) EPC cells seeded on microscopy cover glass in 6-well plates were transfected with 2 μg EGFP-NS79 or EGFP-VP3 and 2 μg mCherry-gcMAVS (G and I), and mCherry-gcTBK1 (H and J). After 24 h, the cells were fixed and subjected to confocal microscopy analysis. Green signals represent overexpressed NS79 or VP3, red signals represent overexpressed gcMAVS or gcTBK1, and blue staining indicates the nucleus region. (original magnification, 63× oil immersion objective). Scale bar, 10 μm. All experiments were repeated at least three times, with similar results.

To further explore the functions of NS79 and VP3, their subcellular locations were monitored in EPC cells. We co-transfected mCherry-gcMAVS or mCherry-gcTBK1 with EGFP-NS79 or EGFP-VP3. Red signals from gcMAVS and gcTBK1 were observed in the cytosol and almost overlapped with the green signals from NS79 and VP3 (Fig 3G-3J). These data suggested that GCRV NS79 and VP3 were colocalized with gcMAVS and gcTBK1 in the cytosol.

### GCRV NS79 is phosphorylated by the gcTBK1 N terminus

To further probe the regulatory mechanism of NS79 on the RLR axis, we analyzed the effect of NS79 on RLR molecules at the protein level. gcMAVS-, gcTBK1-, gcMITA-, gcIRF3-, and gcIRF7-Myc expression vectors were co-transfected with HA-NS79 or an empty vector. As shown in Fig 4A, NS79 had no apparent effects on the RLR factors at the protein level. However, when Myc-NS79 was co-transfected with Flag-gcTBK1, shift bands with higher molecular weights were observed. One possible reason for this observation is that NS79 can be phosphorylated by gcTBK1. To confirm this hypothesis, a dephosphorylation assay was performed *in vitro*. As expected, the shift bands partially disappeared after treatment with calf intestinal phosphatase (CIP), indicating that GCRV NS79 can be phosphorylated by gcTBK1 (Fig 4B). Given that the N-terminal domain is the functional kinase domain for TBK1, the truncated mutant of gcTBK1 was constructed to identify the functional domain on NS79. As shown in Fig 4C, compared to the abundant NS79 phosphorylation found in the wild-type gcTBK1 group, gcTBK1-ΔN (lacking the N terminus) failed to phosphorylate NS79. There results indicate that GCRV NS79 can be phosphorylated by gcTBK1 and that the gcTBK1 N-terminal kinase domain is indispensable.

**Fig 4.**
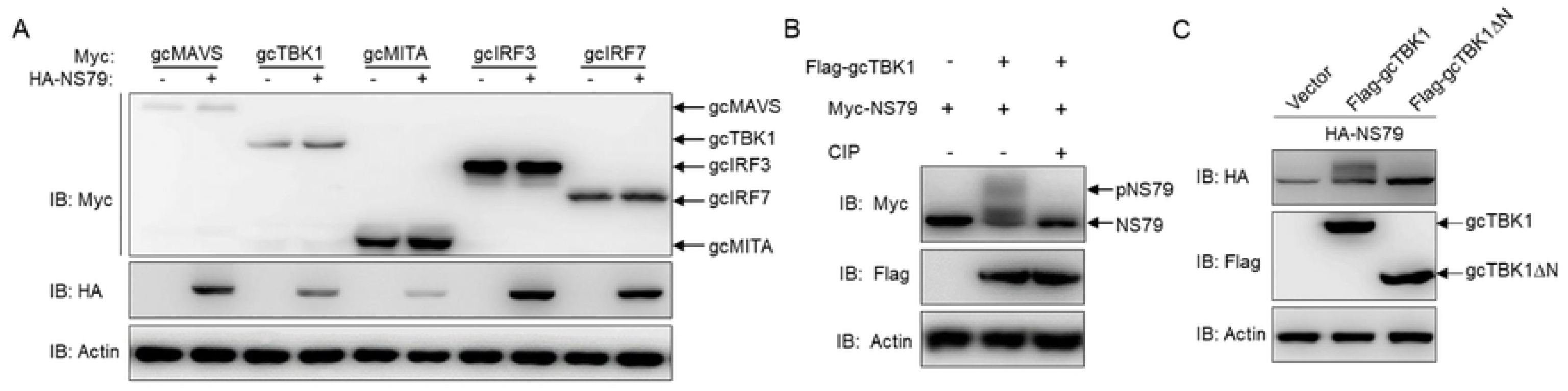
The N terminal of gcTBK1 is essential for the phosphorylation of NS79. (A) NS79 has no effect on the exogenous gcRLR factors. EPC cells were seeded in 6-well plates overnight and transfected with the indicated plasmids (1 μg each) for 24 h. The cell lysates were subjected to IB with anti-Myc, anti-HA, and anti-β-actin Abs. (B) gcTBK1 mediates the phosphorylation of NS79. HEK 293T cells were seeded into 6-well plates overnight and transfected with the indicated plasmids (1 μg each) for 24 h. The cell lysates (100 μg) were treated with or without CIP (10 U) for 40 min at 37°C. Then the lysates were detected by IB with anti-Myc, anti-Flag, and anti-β-actin Abs. (C) gcTBK1-ΔN is indispensable for the phosphorylation of NS79. HEK 293T cells were seeded into 6-well plates overnight and transfected with the indicated plasmids (1 μg each) for 24 h. The cell lysates were subjected to IB with anti-HA, anti-Flag and anti-β-actin Abs. All experiments were repeated at least three times, with similar results.

### NS79 decreases gcTBK1-mediated phosphorylation of gcIRF3

To further determine the biological effect of NS79 on gcTBK1-mediated signaling responses, the functions of gcTBK1 were investigated. As shown in Fig 5A-5C, co-transfection with Flag-gcTBK1 caused a shift of gcMITA, gcIRF3, or gcIRF7 to higher-molecular-weight bands. Subsequently, after the cell lysates were incubated with CIP, the shift bands disappeared, indicating that gcMITA, gcIRF3, and gcIRF7 are also phosphorylated by gcTBK1 in *grass carp*. Furthermore, the truncated mutant of gcTBK1 was used to characterize the functional kinase domain of gcTBK1. Compared with the wild-type gcTBK1, gcTBK1-ΔN was unable to phosphorylate gcMITA, gcIRF3, or gcIRF7 (Fig 5D-5F). These data suggest that the N-terminal kinase domain is also essential for gcMITA, gcIRF3, and gcIRF7 phosphorylation.

**Fig 5.**
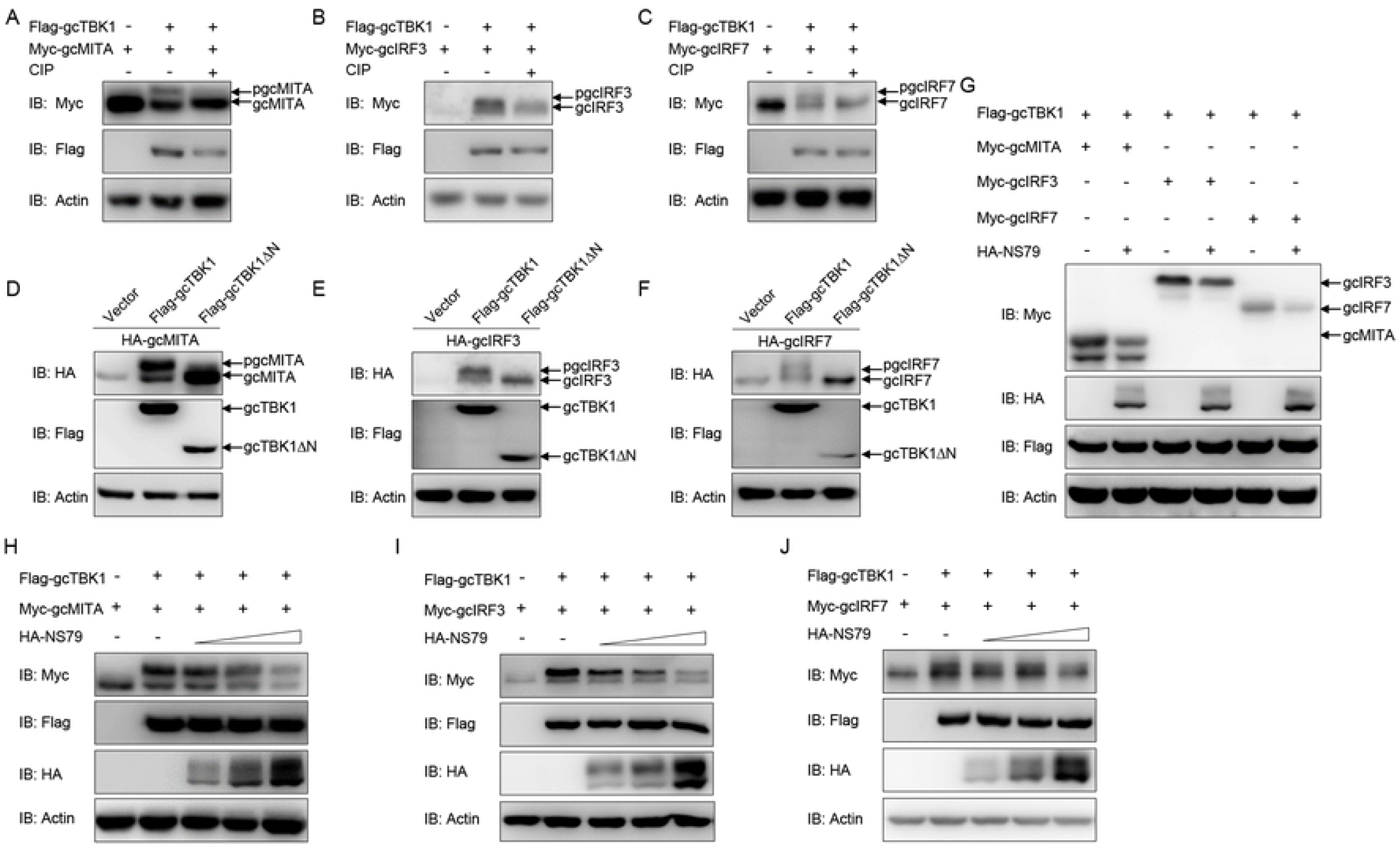
NS79 decreases gcTBK1-mediated phosphorylation of gcMITA, gcIRF3, and gcIRF7. (A-C) gcTBK1 mediates the phosphorylation of gcMITA, gcIRF3 and gcIRF7. HEK 293T cells were seeded into 6-well plates overnight and transfected with the indicated plasmids (1 μg each) for 24 h. The cell lysates (100 μg) were treated with or without CIP (10 U) for 40 min at 37°C. The lysates were then detected by IB with anti-Myc, anti-Flag, and anti-β-actin Abs. (D-F) gcTBK1-ΔN is essential for the phosphorylation of gcMITA, gcIRF3 and gcIRF7. HEK 293T cells were seeded into 6-well plates overnight and transfected with the indicated plasmids (1 μg each) for 24 h. The cell lysates were subjected to IB with anti-HA, anti-Flag and anti-β-actin Abs. (G-J) NS79 decreases gcTBK1-mediated phosphorylation of gcMITA, gcIRF3 and gcIRF7 in a dose-dependent manner. HEK 293T cells were seeded in 6-well plates overnight and transfected with 1 μg Flag-gcTBK1 and 1 μg empty vector or HA-NS79 (G) or HA-NS79 (0.25, 0.5, or 1.0 μg) (H-J), together with 1 μg Myc-gcMITA, Myc-gcIRF3, or Myc-gcIRF7 for 24 h. Then the lysates were subjected to IB with anti-Myc, anti-HA, anti-Flag and anti-β-actin Abs. One representative experiment of at least three independent experiments is shown, and each was done in triplicate. All experiments were repeated at least three times, with similar results.

Next, we wondered whether NS79 affects the gcTBK1-induced phosphorylation of gcMITA, gcIRF3, and gcIRF7. As shown in Fig 5G and 5H, the bands of gcMITA and NS79 exhibited higher mobility when the cells were co-transfected with Flag-gcTBK1, but the phosphorylated gcMITA was gradually reduced by increasing amounts of NS79. Similarly, the gcTBK1-mediated phosphorylation of gcIRF3 and gcIRF7 was also reduced by exogenous expression of NS79 in a dose-dependent manner (Fig 5I and 5J). In conclusion, these results indicate that the GCRV NS79 reduces the gcTBK1-triggered phosphorylation of gcMITA, gcIRF3, and gcIRF7 by being competitively phosphorylated by gcTBK1.

### GCRV VP3 mediates autophagosome-dependent degradation of gcMAVS

The data presented above suggest that NS79 decreases gcTBK1-mediated phosphorylation of IRF3. We next investigated the signaling molecule targeted by VP3. We co-expressed different signaling molecules with HA-VP3 in EPC cells and found that the abundance of gcMAVS was substantially decreased with the overexpression of VP3, but the contents of other RLR factors hardly changed (Fig 6A). In addition, the exogenous gcMAVS was further reduced with overexpressed VP3 in a dose-dependent manner (Fig 6B). Protein degradation is one of the main mechanisms involved in modulating protein functions in biological processes. In general, there are three systems for protein degradation: the ubiquitin-proteasome, autophagosome, and lysosomal pathways. To further investigate the degradation system for gcMAVS, the cells were treated with the indicated inhibitors. The VP3-induced degradation of gcMAVS was completely blocked by the autophagosome inhibitor 3-MA, but not MG132 and NH_4_Cl, the inhibitors of the proteasome and lysosomal pathways, respectively (Fig 6C-6E). Because gcMAVS was degraded by VP3, we speculated that the phosphorylation of gcIRF3 might be impaired. As shown in Fig 6F, the phosphorylation of gcIRF3 mediated by gcMAVS was decreased by overexpression of VP3. Collectively, these data suggest that VP3 degrades gcMAVS in an autophagosome-dependent pathway, which results in the decrease of gcIRF3 phosphorylation.

**Fig 6.**
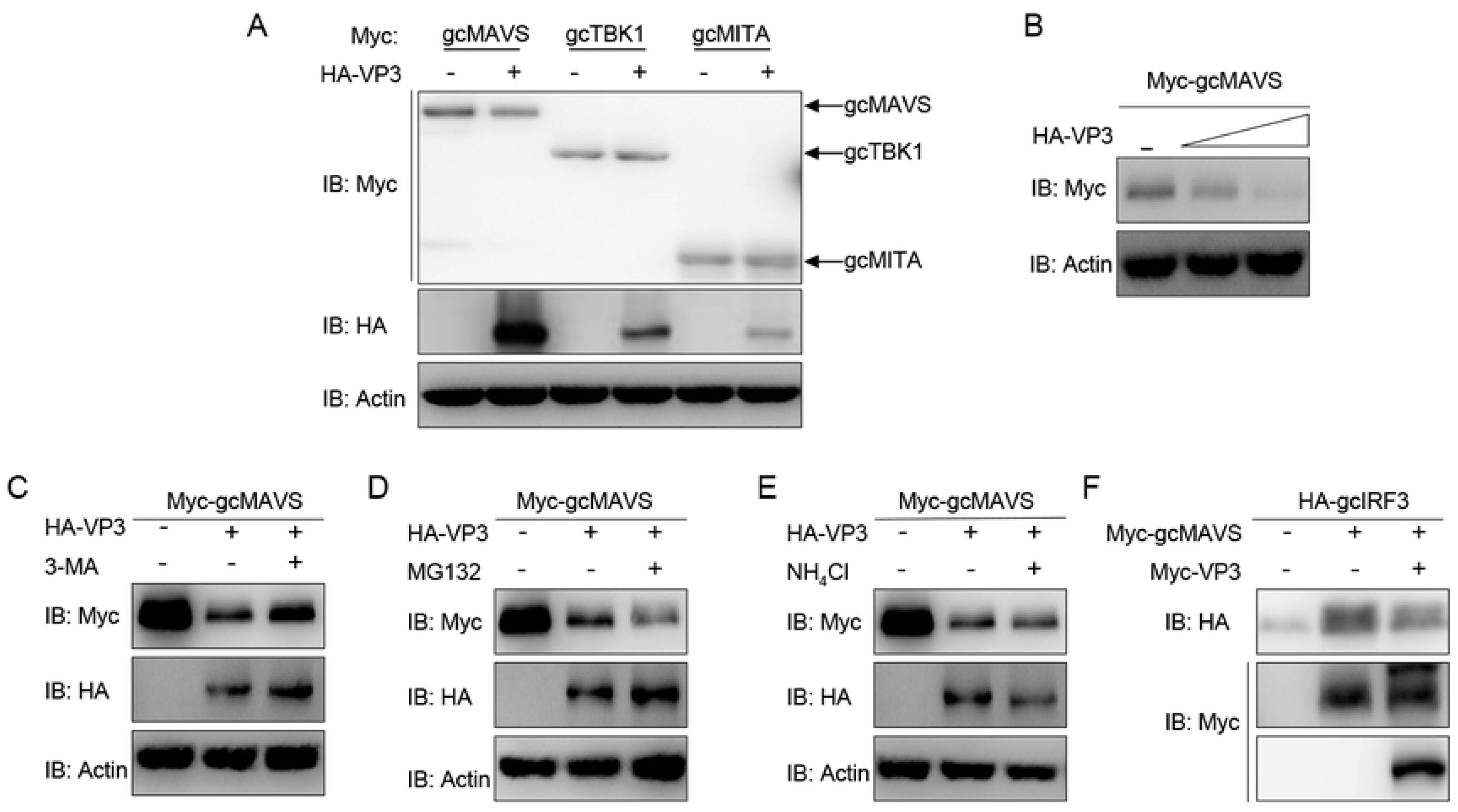
GCRV VP3 degrades gcMAVS through autophagosome pathway. (A) EPC cells were seeded in 6-well plates overnight and transfected the indicated plasmids (1 μg each) for 24 h. The cell lysates were then analyzed by IB with anti-HA, anti-Myc, and anti-β-actin Abs, respectively. (B) Overexpression of the VP3 degrades gcMAVS in a dose-dependent manner. EPC cells were seeded in 6-well plates overnight and co-transfected with 1 μg Myc-gcMAVS and 1 μg empty vector or HA-VP3 (0.5 or 1.0 μg) for 24 h. Then the lysates were subjected to IB with anti-Myc, anti-HA, anti-Flag and anti-β-actin Abs. (C-E) Effects of inhibitors on NS79-mediated degradation of gcMAVS. EPC cells were seeded in 6-well plates overnight and co-transfected with 1 μg Myc-gcMAVS and 1 μg empty vector or HA-VP3 for 18 h, and then treated with DMSO, MG132, 3-MA or NH_4_Cl for 8 h. The cell lysates were subjected to IB with anti-Myc, anti-HA, and anti-β-actin Abs. (F) VP3 decreases gcMAVS-mediated phosphorylation of gcIRF3. HEK 293T cells were seeded in 6-well plates overnight and transfected with 1 μg Myc-gcMAVS and 1 μg empty vector or Myc-VP3, together with 1 μg HA-gcIRF3 for 24 h. The cell lysates were subjected to IB with anti-Myc, anti-HA, and anti-β-actin Abs. All experiments were repeated at least three times, with similar results.

### GCRV NS79 and VP3 attenuate the cellular antiviral response

To ascertain whether GCRV NS79 or VP3 interferes with the cellular IFN response to facilitate virus replication, EPC cells were transfected with NS79 or VP3 and infected with SVCV. As shown in Fig 7A, a stronger CPE was observed in the NS79 and VP3 group at 2 d post-infection. These results were confirmed by the titers of SVCV, which significantly increased 80-fold and 300-fold respectively in the NS79 and VP3-overexpressing cells compared with the control cells (Fig 7B). In addition, qPCR analysis demonstrated that the overexpression of NS79 or VP3 blocked the SVCV-induced expression of *ifn* and *vig1* (Fig 7C and 7D). These data indicate that GCRV NS79 and VP3 suppress the cellular IFN response and facilitate SVCV proliferation.

**Fig 7.**
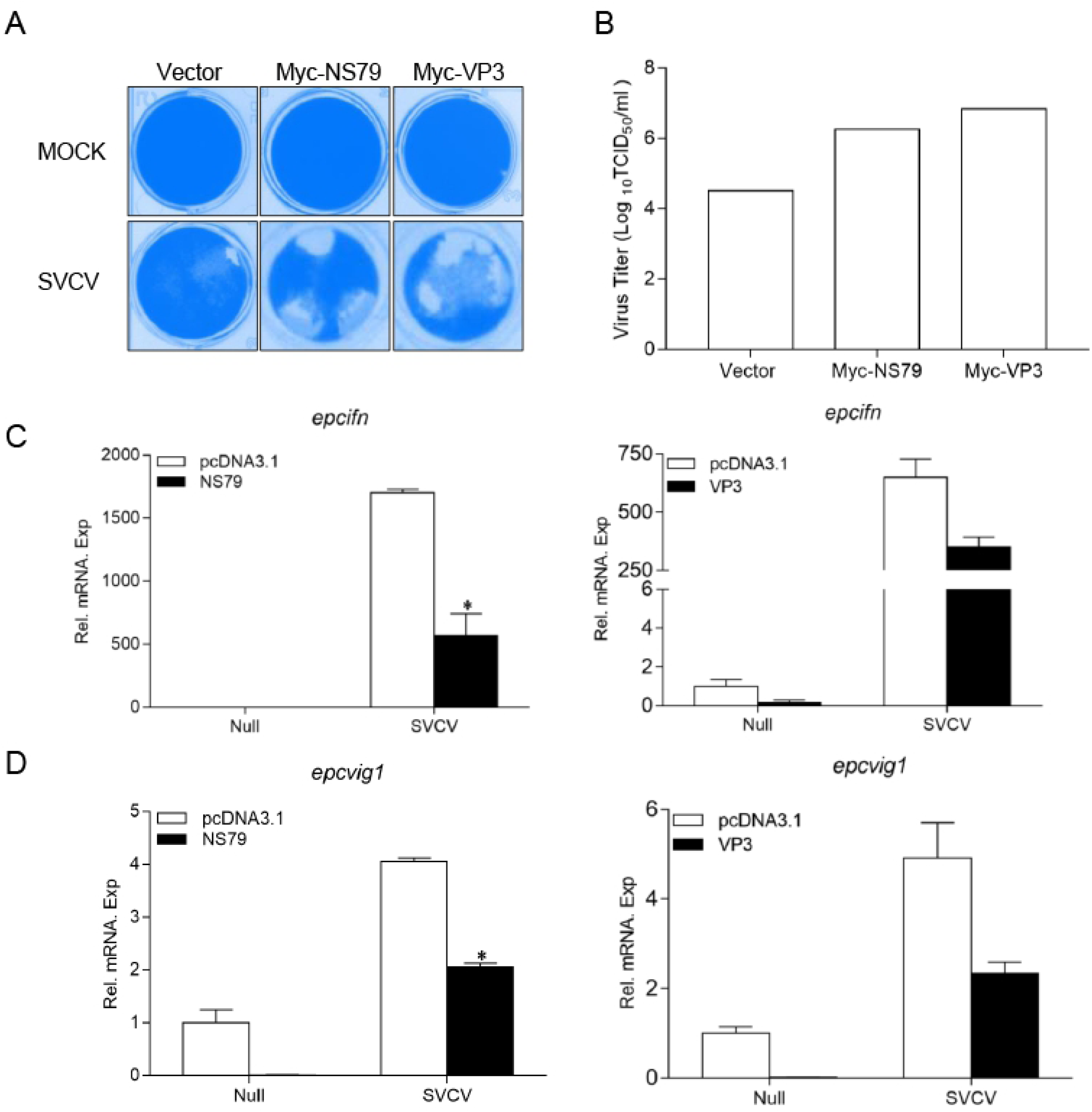
Overexpression of GCRV NS79 or VP3 dampens the cellular IFN responses. (A and B) Increase of virus replication by overexpression of NS79 or VP3. EPC cells were seeded in 24-well plates overnight and transfected with indicated plasmids for 24 h, then the cells were infected with SVCV (MOI = 0.001) for 48 h. (A) Then, the cells were fixed with 4% PFA and stained with 1% crystal violet. (B) Culture supernatants from the cells infected with SVCV were collected, and the viral titer was measured by TCID_50_ assays. (C and D) overexpression of NS79 or VP3 inhibits SVCV-induced up-regulation of *epcifn* (C) and *epcvig1* (D). EPC cells seeded in 6-well plates overnight were transfected with 2 μg empty vector or pcDNA3.1-NS79, or pcDNA3.1-VP3, at 24 h post-transfection, the cells were infected with SVCV (MOI = 1) for 24 h. The total RNAs were extracted to examine the mRNA levels of cellular *epcifn* and *epcvig1*. The relative transcriptional levels were normalized to the transcriptional level of the *β-actin* gene and were represented as fold induction relative to the transcriptional level in the control cells, which was set to 1. Data were expressed as mean ± SEM, *n* = 3. Asterisks indicate significant differences from control values (*, *p* < 0.05). Experiments were repeated for at least three times with similar results.

## Discussion

In a water living environment, aquatic viruses can spread more easily and cause higher mortality than land-based viruses. In previous studies, our lab has reported that GCRV VP41 protein (encoded by the S8 segment) as well as SVCV N and P proteins antagonize fish RLR factors to reduce host IFN production [41-43], highlighting the evasion mechanisms used by aquatic viruses. To date, precise information regarding both the fish IFN response to viruses and the pathogenesis of aquatic viruses is rare; thus, the immune evasion mechanisms of aquatic viruses targeting the host IFN system are unclear. Here, we report that all GCRV viral proteins can interact with host RLR factors, specifically inhibiting the MAVS-mediated production of IFNs. Furthermore, we found that NS79 reduces gcMITA phosphorylation by acting as a decoy substrate of gcTBK1 while VP3 degrades gcMAVS in an autophagosome-dependent manner, ultimately inhibiting IFN production and facilitating virus replication. This finding enhances the understanding of the immune evasion mechanisms of GCRV.

It is obviously rare for all proteins in one virus to interfere with the TBK1 kinase, hence we presume the observed phenomenon has two causes: (1) TBK1 is crucial for fish IFN production. More type I IFN members are found in fish than in mammals (which possess only IFNα and IFNβ), and the regulation patterns are more complicated. TBK1 is the upstream kinase of IRF3, IRF7, and even IRF6 (positive regulator of fish IFNs), displaying a powerful capacity to activate IFNs [44]; therefore, targeting TBK1 is more efficient for a virus than targeting each IRF. (2) TBK1 has multiple functions in hosts. Besides activating IFN transcription, TBK1 participates in several host life processes such as cellular transformation, autophagy, antibacterial response, and oncogenesis [25, 45]. For example, autophagy is a conserved process in eukaryotic cells and plays a crucial role in the eukaryotic defense against pathogens. TBK1 interacts with and phosphorylates optineurin (OPTN, a key component of pathogen-induced autophagy), leading to the elimination of pathogens by xenophagy [46]. For viral infection, viruses also need to control host resources for replication and proliferation in addition to combating the host IFN response. Thus, choosing TBK1 as the target is a highly effective way for GCRV to proliferate in host cells. The cell physiology mediated by GCRV-regulated TBK1 should be elucidated with further studies on the biological function of fish TBK1.

TBK1 is a pivotal protein kinase that is utilized by viruses [47]. In the viral lifecycle, the phosphorylation of viral proteins has been identified as indispensable, and several studies have suggested that viral proteins are not active until phosphorylated by cellular kinase. The phosphorylation of NS1 protein from *Periplaneta fuliginosa* densovirus (PfDNV) triggers the activation of viral genome replication and transcription [48]. IE63 from vesicular stomatitis virus (VSV) is phosphorylated by host cellular cyclin-dependent kinase (CDK) 1 and CDK2, then translocates from the nucleus to the cytoplasm [49]. Human immunodeficiency virus 1 (HIV-1) Gag and Vpr proteins improve their assembly into viral particles after phosphorylation by host atypical protein kinase C (aPKC) [50]. The phosphorylation of viral proteins has also been observed in aquatic viruses. For example, SVCV P protein can be phosphorylated by TBK1, which leads to the decline of IRF3 phosphorylation and IFN production [43]. However, the function of phosphorylated P protein in SVCV proliferation is still unknown. In this study, the NS79 protein encoded by the GCRV S4 segment was also phosphorylated by TBK1. As S4 encodes a non-structural protein and is possibly involved in the formation of viral inclusion bodies, with the protein encoded by GCRV S9, the function of phosphorylated NS79 might be indispensable for viral assembly.

Conversely, the physical binding of TBK1 with viral proteins might inhibit the host’s antiviral response. Since TBK1 is considered to have a pivotal antiviral role in phosphorylating IRF3 to activate IFN transcription, the significant disruption of the signaling transduction of TBK1 by viruses will blunt the host IFN production [22]. Meanwhile, the viral protein(s) that interact with host TBK1 may also be crucial for the viral life cycle, and such neutralization will disrupt the normal transcription, translation, and proliferation of viruses. For example, the borna disease virus (BDV) P protein associates with TBK1 and inhibits its kinase activity to promote viral evasion [51]. The rhabdovirus P protein interacts with the L protein binding with the viral template in the transcription process, which facilitates the N protein staying in a soluble, encapsidation-competent form that is associated with viral RNA to form the nucleocapsid during viral assembly [52, 53]. The P protein amount is reduced after reacting with the host TBK1; a lower concentration of the P protein should reduce the normal viral transcription level. The outcome of the combat between virus and host might be determined by the amount and role of the proteins that participate on both sides, as well as the reaction efficiency; however, the exact mechanisms need to be further clarified. The current study identified a novel phenomenon of aquatic virus GCRV countering the host IFN response. In the most common and highest mortality genotype of GCRV, GCRV II, all the viral proteins encoded by the segments reduce the host IFN transcription by interacting with TBK1. We hope our findings provide a base for further study of GCRV evasion mechanisms related to TBK1.

The critical role of MAVS in the production of IFN and other proinflammatory cytokines predisposes it to being a target of many viruses [54]. In long-term coexistence of virus and host, viruses have evolved various strategies to suppress MAVS-mediated signaling. One of the most common mechanisms is the cleavage of MAVS, resulting in the dislocation of MAVS from the mitochondria, thus preventing IFN induction. For instance, the HCV protease NS3/4A and Enterovirus 71 Protease 2A^pro^ cleave MAVS to block signaling transduction [55, 56]. Besides the cleavage of MAVS, several viruses choose to degrade MAVS. For example, hepatitis B virus X protein (HBX) interacts with MAVS, inducing the degradation of MAVS through Lys^136^ ubiquitin in MAVS protein, thus inhibiting IFN expression [57]. Our finding that GCRV VP3 interacts with and degrades MAVS in an autophagosome -dependent manner provides new insight into how virus-derived proteins and MAVS can interact.

Overall, we showed that GCRV attenuates host immune signaling mediated by two potent antiviral adapter molecules, MAVS and TBK1. This is achieved using two distinct methods to reduce MAVS and TBK1 signaling, namely VP3 triggering the degradation of MAVS and NS79 serving as a substrate of TBK1 to reduce IRF3 phosphorylation. Our findings therefore revealed new mechanisms of GCRV-mediated evasion of the host’s innate immunity.

## Materials and methods

### Cells and viruses

Human embryonic kidney (HEK) 293T cells were provided by Dr. Xing Liu (Institute of Hydrobiology, Chinese Academy of Sciences) and were grown at 37°C in 5% CO_2_ in Dulbecco’s modified Eagle’s medium (DMEM; Invitrogen) supplemented with 10% fetal bovine serum (FBS, Invitrogen). Grass carp ovary (GCO) cells and Epithelioma papulosum cyprini (EPC) cells were obtained from China Center for Type Culture Collection (CCTCC) and were maintained at 28°C in 5% CO_2_ in medium 199 (Invitrogen) supplemented with 10% FBS. GCRV (strain 106, group II) was a gift from Lingbing Zeng (Yangtze River Fisheries Research Institute, Chinese Academy of Fishery Sciences). Because group II GVRV cannot cause a cytopathic effect (CPE) but can propagate in GCO cells, the cultured media with GCO cells infected with group II GCRV for 8 days were harvested and stored at −80°C until used. Spring viremia of carp virus (SVCV), a negative single-stranded RNA (ssRNA) virus, was propagated in EPC cells until CPE was observed; then the culture medium with cells was harvested and stored at −80°C until used.

### Plasmid construction and reagents

The open reading frame (ORF) of GCRV S1-S11 (KC201166.1, KC201167.1, KC201168.1, KC201169.1, KC201170.1, KC201171.1, KC201172.1, KC201173.1, KC201174.1, KC201175.1, KC201176.1) were generated by PCR and then cloned into pcDNA3.1(+) (Invitrogen), pCMV-Myc (Clontech), or pCMV-HA vectors (Clontech), respectively. The ORFs of gcRIG-I (GQ478334.2), gcMAVS (KF366908.1), gcTBK1 (JN704345.1), gcMITA (JN786909.1), gcIRF3 (KT347289.1), and gcIRF7 (KY613780.1) were also subcloned into pcDNA3.1(+), pCMV-Myc, pCMV-HA, and pCMV-Tag2C vectors, respectively. For subcellular localization, the ORFs of VP3 and NS79 were inserted into pEGFP-N3 vector (Clontech), respectively. The ORFs of gcMAVS and gcTBK1 were also inserted into pCS2-mCherry vector (Clontech). The expression plasmids for Flag/pcDNA3.1-DrMAVS, Flag/pcDNA3.1-DrTBK1, Flag-DrMITA, Flag-DrIRF3, and Flag-DrIRF7 were described previously [43]. For promoter activity analysis, gcIFN1/gcIFN2/gcIFN3/gcIFN4pro-Luc construct were generated by insertion of corresponding 5′-flanking regulatory region of gcIFN1 promoter (GU139255.1), gcIFN2 promoter (KY613781.1), gcIFN3 promoter (KY613782.1), or gcIFN4 promoter (KY613783.1) into pGL3-basic luciferase reporter vector (Promega, Madison, WI), respectively. The DrIFNφ1pro-Luc and ISRE-Luc plasmids in the pGL3-basic luciferase reporter vector (Promega) were constructed as described previously [42]. The Renilla luciferase internal control vector (pRL-TK) was purchased from Promega. The primers including the restriction enzyme cutting sites used for plasmid construction are listed in S1 Table. All constructs were confirmed by DNA sequencing.

### Luciferase activity assay

EPC cells or GCO cells were seeded in 24-well plates overnight and co-transfected with the indicated luciferase reporter plasmid and overexpression plasmid. The empty vector pcDNA3.1(+) was used to ensure equivalent amounts of total DNA in each well. Transfection of poly I:C was performed at 24 h before cell harvest. At 48 h post-transfection, the cells were washed with phosphate-buffered saline (PBS) and lysed for measuring luciferase activity by the Dual-Luciferase Reporter Assay System (Promega) according to the manufacturer’s instructions. Firefly luciferase activity was normalized on the basis of Renilla luciferase activity. At least three independent experiments were performed and the data from one representative experiment were conducted for statistical analysis.

### RNA extraction, reverse transcription, and qPCR

Total RNA was extracted by the TRIzol reagent (Invitrogen). First-strand cDNA was synthesized by using a GoScript reverse transcription system (Promega) according to the manufacturer’s instructions. qPCR was performed with Fast SYBR green PCR master mix (Bio-Rad) on the CFX96 real-time system (Bio-Rad). PCR conditions were as follows: 95°C for 5 min and then 40 cycles of 95°C for 20 s, 60°C for 20 s, and 72°C for 20 s. All primers used for qPCR are shown in S1 Table, and the β-actin gene was used as an internal control. The relative fold changes were calculated by comparison to the corresponding controls using the 2-ΔΔCt method. At least three independent experiments were performed and the data from one representative experiment were conducted for statistical analysis.

### Transient transfection and virus infection

Transient transfections were performed in EPC cells seeded in 6-well or 24-well plates by using X-tremeGENE HP DNA Transfection Reagent (Roche) according to the manufacturer’s protocol. For the antiviral assay using 24-well plates, EPC cells were transfected with 0.5 μg pcDNA3.1-VP3/NS79 or the empty vector. At 24 h post-transfection, cells were infected with SVCV at a multiplicity of infection (MOI = 0.001). After 2 or 3 d, supernatant aliquots were harvested for detection of virus titers, the cell monolayers were fixed by 4% paraformaldehyde (PFA) and stained with 1% crystal violet for visualizing CPE. For virus titration, 200 μl of culture medium were collected at 48 h post-infection, and used for plaque assay. The supernatants were subjected to 3-fold serial dilutions and then added (100 μl) onto a monolayer of EPC cells cultured in a 96-well plate. After 48 or 72 h, the medium was removed and the cells were washed with PBS, fixed by 4% PFA and stained with 1% crystal violet. The virus titer was expressed as 50% tissue culture infective dose (TCID_50_/ml). Results are representative of three independent experiments.

### Co-immunoprecipitation (Co-IP) assay

The HEK 293T cells seeded in 10 cm^2^ dishes overnight were transfected with a total of 10 μg of the plasmids indicated on the figures. At 24 h post-transfection, the medium was removed carefully, and the cell monolayer was washed twice with 10 ml ice-cold PBS. Then the cells were lysed in 1 ml of radioimmunoprecipitation (RIPA) lysis buffer [1% NP-40, 50 mM Tris-HCl, pH 7.5, 150 mM NaCl, 1 mM EDTA, 1 mM NaF, 1 mM sodium orthovanadate (Na_3_VO_4_), 1 mM phenyl-methylsulfonyl fluoride (PMSF), 0.25% sodium deoxycholate] containing protease inhibitor cocktail (Sigma-Aldrich) at 4°C for 1 h on a rocker platform. The cellular debris was removed by centrifugation at 12,000 × *g* for 15 min at 4°C. The supernatant was transferred to a fresh tube and incubated with 30 µl anti-HA-agarose beads or anti-Flag/Myc affinity gel (Sigma-Aldrich) overnight at 4°C with constant agitation. These samples were further analyzed by immunoblotting (IB). Immunoprecipitated proteins were collected by centrifugation at 5000 × *g* for 1 min at 4°C, washed three times with lysis buffer and resuspended in 50 μl 2 × SDS sample buffer. The immunoprecipitates and whole cell lysates were analyzed by IB with the indicated antibodies (Abs).

### Immunoblot analysis

Immunoprecipitates or whole cell lysates were separated by 10% SDS-PAGE and transferred to polyvinylidene difluoride (PVDF) membrane (Bio-Rad). The membranes were blocked for 1 h at room temperature in TBST buffer (25 mM Tris-HCl, 150 mM NaCl, 0.1% Tween 20, pH 7.5) containing 5% nonfat dry milk, probed with the indicated primary Abs at an appropriate dilution overnight at 4°C, washed three times with TBST, and then incubated with secondary Abs for 1 h at room temperature. After three additional washes with TBST, the membranes were stained with the Immobilon Western chemiluminescent horseradish peroxidase (HRP) substrate (Millipore) and detected by using an ImageQuant LAS 4000 system (GE Healthcare). Abs were diluted as follows: anti-*β-actin* (Cell Signaling Technology) at 1:1,000, anti-Flag/HA (Sigma-Aldrich) at 1:3,000, anti-Myc (Santa Cruz Biotechnology) at 1:2,000, and HRP-conjugated anti-mouse IgG (Thermo Scientific) at 1:5,000. Results are representative of three independent experiments.

### In vitro protein dephosphorylation assay

Transfected HEK 293T cells were lysed as described above, except that the phosphatase inhibitors (Na_3_VO_4_ and EDTA) were omitted from the lysis buffer. Protein dephosphorylation was carried out in 100 μl reaction mixtures consisting of 100 μg of cell protein and 10 units (U) of CIP (Sigma-Aldrich). The reaction mixtures were incubated at 37°C for 40 min, followed by immunoblot analysis.

### Fluorescent microscopy

EPC cells were plated onto coverslips in 6-well plates and transfected with the plasmids indicated on the figures for 24 h. Then the cells were washed twice with PBS and fixed with 4% PFA for 1 h. After being washed three times with PBS, the cells were stained with 1 µg/ml 4′, 6-diamidino-2-phenylindole (DAPI; Beyotime) for 15 min in the dark at room temperature. Finally, the coverslips were washed and observed with a confocal microscope under a 63× oil immersion objective (SP8; Leica).

### Statistics analysis

Luciferase and qPCR assay data are expressed as the mean ± standard error of the mean (SEM). Error bars indicate the SEM (*n* = 3, biologically independent samples). The *p* values were calculated by one-way analysis of variance (ANOVA) with Dunnett’s post hoc test (SPSS Statistics, version 19; IBM). A *p* value < 0.05 was considered statistically significant.

## Acknowledgments

We thank Dr. Fang Zhou (Institute of Hydrobiology, Chinese Academy of Sciences) for assistance with confocal microscopy analysis.

## Supporting Information legends

**S1 Fig. The segments of GCRV suppresses the DrMAVS-mediated activation of IFN1**. (A and B) The segments of GCRV inhibits DrIFNφ1 (A) and ISRE (B) activation mediated by DrMAVS. GCO cells were seeded in 24-well plates and transfected with pcDNA3.1-DrMAVS and empty vector or pcDNA3.1-S1, or pcDNA3.1-S2, or pcDNA3.1-S3, or pcDNA3.1-S4, or pcDNA3.1-S5, or pcDNA3.1-S6, or pcDNA3.1-S7, or pcDNA3.1-S8, or pcDNA3.1-S9, or pcDNA3.1-S10, or pcDNA3.1-S11, plus DrIFNφ1pro-Luc (A), or ISRE-Luc (B) at the ratio of 1:1:1. pRL-TK was used as a control. At 24 h post-transfection, cells were lysed for luciferase activity detection. (C and D) The segments of GCRV have no apparent effects on DrTBK1-mediated activation of DrIFNφ1 (C) and ISRE reporters (D). GCO cells were seeded in 24-well plates and transfected with pcDNA3.1-gcTBK1 and empty vector or pcDNA3.1-S1, or pcDNA3.1-S2, or pcDNA3.1-S3, or pcDNA3.1-S4, or pcDNA3.1-S5, or pcDNA3.1-S6, or pcDNA3.1-S7, or pcDNA3.1-S8, or pcDNA3.1-S9, or pcDNA3.1-S10, or pcDNA3.1-S11, plus DrIFNφ1pro-Luc (C), or ISRE-Luc (D) at the ratio of 1:1:1. pRL-TK was used as a control. At 24 h post-transfection, cells were lysed for luciferase activity detection. The promoter activity is presented as relative light units normalized to Renilla luciferase activity. Data were expressed as mean ± SEM, n = 3. Asterisks indicate significant differences from control (*, p < 0.05). All experiments were repeated for at least three times with similar result.

**S2 Fig. The segments of GCRV associates with the key molecules of zebrafish RLR signaling pathway**. HEK 293T cells seeded in 10-cm^2^ dishes were transfected with the indicated plasmids (5 μg each). After 24 h, cell lysates were IP with an anti-Flag affinity gel. Then the immunoprecipitates and cell lysates were analyzed by IB with the anti-Myc and anti-Flag Abs, respectively. All experiments were repeated at least three times, with similar results.

